# The mechanical impact of *col11a2* loss on joints; *col11a2* mutant zebrafish show changes to joint development and function, which leads to early onset osteoarthritis

**DOI:** 10.1101/302307

**Authors:** Elizabeth A Lawrence, Erika Kague, Jessye A Aggleton, Robert L Harniman, Karen A Roddy, Chrissy L Hammond

**Affiliations:** School of Physiology, Pharmacology and Neuroscience, University of Bristol BS8 1TD; School of Anthropology and Archaeology, University of Bristol, BS8 1UU; School of Chemistry, University of Bristol, BS8 1TS

**Keywords:** Zebrafish, biomechanics, collagen, cartilage, Stickler syndrome, development

## Abstract

Collagen is the major structural component of cartilage and mutations in the genes encoding Type XI collagen are associated with severe skeletal dysplasias (Fibrochondrogenesis and Stickler syndrome) and early onset osteoarthritis. The impact of the lack of Type XI collagen on cell behaviour and mechanical performance during skeleton development is unknown. We studied a zebrafish mutant for *col11a2* and evaluated cartilage, bone development and mechanical properties to address this. We show that in *col11a2* mutants Type II collagen is made but is prematurely degraded in maturing cartilage and ectopically expressed in the joint. These changes are correlated with increased stiffness of both bone and cartilage; quantified using Atomic Force Microscopy. In the mutants, the skeletal rudiment terminal region in the jaw joint are broader and the interzone smaller. These differences in shape and material properties impact on joint function and mechanical performance, which we modelled using Finite Element Analyses. Finally, we show that *col11a2* heterozygous carriers reach adulthood but show signs of severe early onset osteoarthritis. Taken together our data demonstrate a key role for Type XI collagen in maintaining the properties of cartilage matrix; which when lost leads to alterations to cell behaviour that give rise to joint pathologies.

## Introduction

Articular cartilage is a highly specialised connective tissue, which provides a smooth, lubricated surface for articulation and load transmission with low joint friction. Collagen is a major constituent of cartilage, accounting for around 75% of its dry weight (1). Type II collagen makes up 90-95% percent of the collagen network, while the remaining 5-10% is comprised of other collagens such as Type IX and XI, with studies of chick articular cartilage showing association of these three collagen types in a tight D-periodic array (2,3) These minor collagens help to organise and stabilize the Type II collagen fibril network that, along with proteoglycans, water and other proteins, form a dense extracellular matrix (ECM) in which chondrocytes are dispersed (4). The tight fibrillar structure from collagens and the water content, from interaction with glycosaminoglycans (GAGs), govern the mechanical properties of the cartilage. Type XI collagen belongs to the fibril-forming class of collagens; it is formed as a heterotrimer of three chains each encoded by a different gene: *COL11A1* (5), *COL11A2,* and *COL2A1(6,7).* While the α1 chain of Type XI collagen is expressed in both cartilaginous and ocular tissue, the α2 chain is predominantly expressed in cartilage.

Given the close interaction between Type II and Type XI collagens, mutations that affect either can cause similar destabilization of cartilage organisation, as observed in Marshall Stickler Syndrome. Stickler syndrome, which affects around 1 in 7500 new-borns, encompasses a hereditary group of conditions caused by defective Type II, IX or XI Collagen (8) and is divided in to three phenotypes: depending on the collagen mutation present. Type III is associated with mutations to the Type XI gene *COL11A2* (9) (10). Type III Stickler syndrome is characterised by skeletal, orofacial, and auditory abnormalities including: scoliosis; hearing loss; cleft palate; joint hypermobility; (11), multiple hereditary exostoses (10), and premature osteoarthritis (OA) in 75% of patients before the age of 30 (8). The majority of mutations linked to Stickler syndrome lead to truncated proteins lacking the c-terminal domain of the peptide, disturbing the association of the alpha helices to form procollagens and consequentially the formation of collagen fibrils and fibres (12). Mutations in genes encoding Type XI collagens are also associated with other skeletal dysplasias, including the severe developmental condition Fibrochondrogenesis (13), and Weissenbacher-Zymuller syndrome (9).

Mutant mice for *Col11a1 (Cho-/-),* are neonatally lethal and show decreased limb bone length, cleft palate and short snouts (14), and thicker, less uniform collagen fibrils in the cartilage ECM (15). Type II collagen degradation (16) and early onset OA were reported in *Cho/+* heterozygous mice (17). Additionally, mice haploinsufficient for *Col11a1* display altered susceptibility to load induced damage (18). While *Col11a2* mutant mice have been reported to show hearing loss, their skeletal phenotype has not been described (19). The interaction of Type XI collagen with Type II is important for the maintenance of the spacing and diameter of Type II collagen fibrils (20). As Type II collagen is the major collagen in cartilage, changes to its organisation can impact the mechanical performance of the cartilage. Computational modelling has shown that spacing and interconnectivity between collagen fibrils has a significant effect on the mechanical performance of cartilage (21). Cartilage is an intrinsically mechanically sensitive tissue, and changes to cartilage biomechanical performance have been extensively described during development (22), ageing and disease (23). It is also increasingly well understood that subtle changes to skeletal morphology and joint shape can increase susceptibility to joint conditions such as osteoarthritis later in life (24). A recent large GWAS on hip shape identified *COL11A1* as a contributor to hip shape (25). Joint shapes seen in development and disease have been shown to have significant impact on the biomechanical performance of joints (24,26). What is less clear is the sequence of events within the joint, do changes to shape precede changes to cartilage structure and mechanical performance or vice versa, and what is the relative impact of each change?

Zebrafish are an attractive model for studying the effect of genetic lesions on skeletal development. The larvae are translucent which, twinned with fluorescent reporter transgenic lines, enables dynamic imaging of skeletal cells (27,28) and the development of the zebrafish craniofacial skeleton is well documented (29–31). The zebrafish jaw joint is synovial (32) and requires mechanical input to form normally (33,34) Acute knockdown of *col11a1* in zebrafish using morpholinos has been shown to affect chondrocyte maturation (35), but no stable mutants for *col11a1* or *col11a2* have previously been reported.

Here, we show that larval zebrafish carrying a *col11a2* mutation display a variety of phenotypes including alterations to: joint shape, cartilage composition, cell organisation and the material properties of the cartilage during development. These changes impact on the biomechanical and functional performance of the joint. The mutant fish go on to display phenotypes consistent with Stickler syndrome such as altered face shape and early onset osteoarthritis. Taken together these data suggest that mechanical and cellular changes to the developing skeleton explain the predisposition of people with mutations in Type XI Collagens to early onset osteoarthritis.

## Methods

### Zebrafish husbandry and transgenic lines

Zebrafish were maintained as described previously (36), all experiments were approved by the local ethics committee and performed under a UK Home Office Project Licence. Transgenic lines *Tg(col2a1aBAC:mcherry), Tg(col10a1aBAC:citrine)^hu7050^* (37) and *Tg(smyhc1:GFP)* (38), have been described previously *col11a2^sa18324^* mutant zebrafish was generated by the Zebrafish Mutation Project (Sanger Institute) and acquired from the European Zebrafish Resource Centre (EZRC). It carries a nonsense mutation (C>A base pair change at position 228aa, zv9 chr19: 7834334), leading to a premature stop codon which shortens the polypeptide to about 1/3 of the triple helical domain of the α2 chain of collagen XI.

### DNA extraction and genotyping

Fins were clipped from anaesthetised or fixed zebrafish and incubated in base solution (25 mM NaOH,0.2 mM EDTA) before the addition of neutralisation solution (40 mM Tris HCl, Ph5.0). For genotyping we used KASP (LGC) genotyping or PCR followed by Sanger sequencing *(col11a2* F-GGTGGCCTGATTCTGACCA; *col11a2* R-TATCTCACACCAGGATGCCG). Mutants were identified by C>A base pair change at position 228aa.

### Wholemount immunohistochemistry

Performed as previously described (37). Primary antibodies and dilutions used were: rabbit pAb to collagen II, *(abcam* ab34712), 1:500; mouse pAb to collagen II, *(DHSB* II-II6B3), 1:500, rabbit pAb to collagen I, *(abcam* ab23730), 1:100. Secondary antibodies were Dylight 488 or 550 *(Thermo Scientific)* used at a dilution of 1:500. For imaging, larvae were mounted ventrally in 1% agarose and imaged on a Leica SP5 confocal microscope with a 10x objective.

### 3D render generation, joint measurements, quantification of exostoses and cell circularity

3D volume renders, surface models and measurements were acquired using Amira 6.0 (FEI). Surface models were generated manually by segmenting jaw joints using the segmentation tool. Measurements, as depicted in Figure 1C-D and 3A were taken using the 3D perspective measurement tool. To better visualise exostoses, 3D volume renders were created, and the grayscale range of colour applied. Exostoses were quantified in each lower jaw element from single confocal image stacks in ImageJ (39) using the multi-point tool. Cell circularity was measured from confocal image stacks of Type II collagen immunostained zebrafish larvae at 5dpf. The freehand selection tool in ImageJ was used to outline chondrocytes in three distinct jaw regions (shown in Figure 2H) and the measure function was used to analyse the circularity of each cell. This was done for 10 cells in each region, in 3 wt and 3 *col11a2* mutant zebrafish.

**Figure 1:**
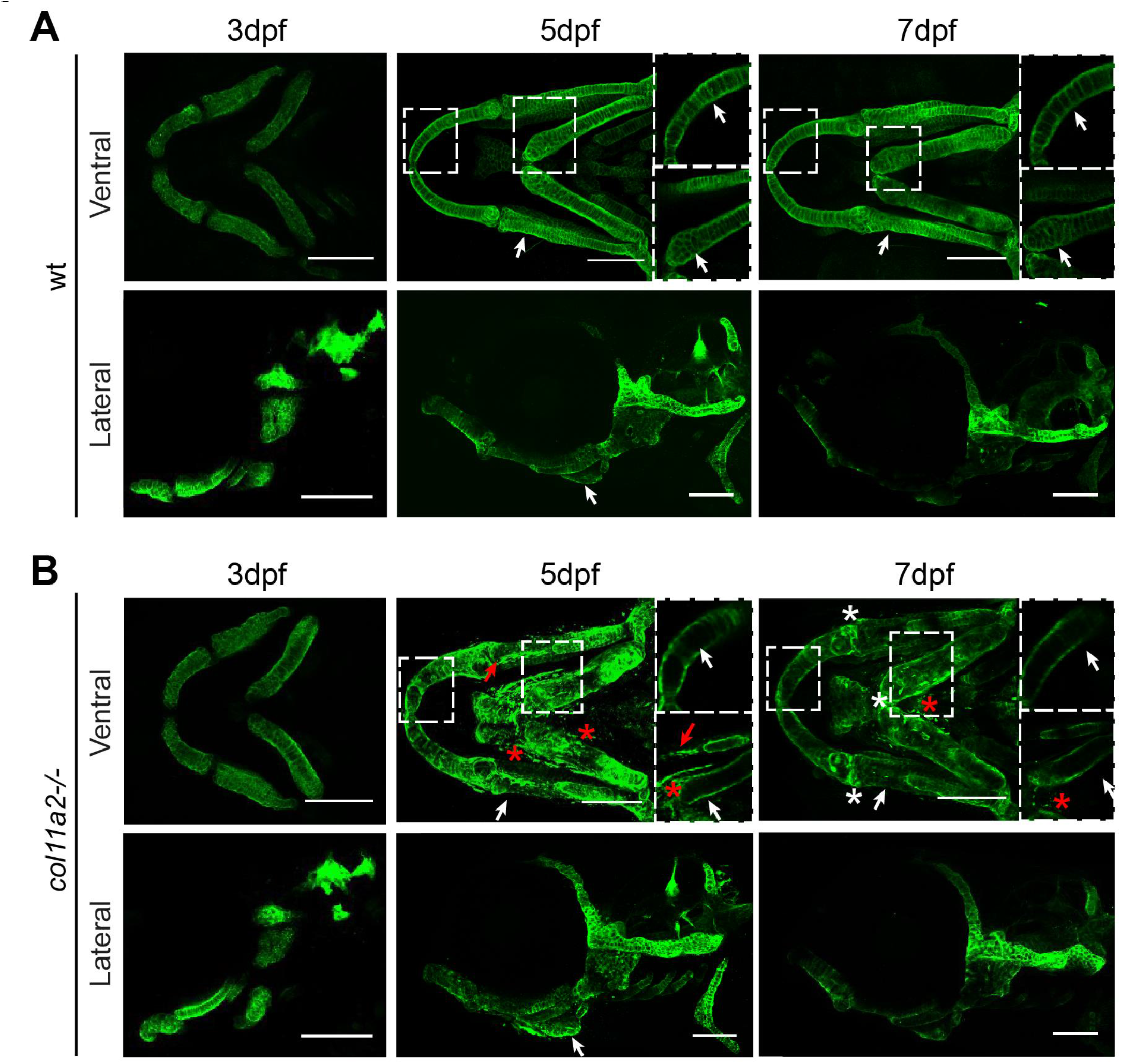
*col11a2* zebrafish mutant larvae show progressively altered Type II collagen protein localisation in jaw cartilage. A, B) Maximum projection of ventral and lateral confocal image stacks from wild type (wt) (A) and homozygous mutant *(col11a2-/-)* (B) larvae immunostained for Type II collagen at three time points (3, 5 and 7dpf). White arrows indicate areas of change in Type II collagen distribution in the extracellular matrix. Dashed insets show single stack images of regions with reduced deposition (white asterisks represent areas where Type II collagen is maintained in mutant fish and red asterisks show fragments of Type II collagen positive material outside the main cartilage elements). Red arrows show interoperculomandibular (IOM) ligament. Scale bar = 100μm.

### Live imaging of transgenic fish

Live larvae at 5dpf were anaesthetised in 0.1mg/ml MS222 and mounted ventrally in

0.3% agarose with tricaine prior to being imaged on a Leica SP5II confocal microscope with a 10x objective. The number of slow muscle fibres and *col10a1a* expressing cells were quantified manually in ImageJ from confocal images of double transgenic *Tg(smyhc1:GFP);(Col2a1aBAC:mcherry)* and *Tg(col10a1aBAC:citrine); (col2a1aBAC:mCherry)* zebrafish at 5dpf, respectively.

### Alcian blue and alizarin red staining

5dpf and 7dpf wt and *col11a2* mutant larvae were stained following a previously described protocol (40) and imaged on a Leica MZ10F stereo microscope prior to genotyping.

### *In situ* hybridisation

*In situ* hybridisation was performed as described (41) using a previously described *col11a2* probe (42). Larvae were imaged on GXM-L3200 B upright microscope.

### Nanoscale surface morphology and Young’s Moduli

Atomic Force Microscopy (AFM) was performed on adult (1 year) bone and larval (7dpf) cartilage from wt and *col11a2* mutant fish. A multi-mode VIII AFM with Nanoscope V controller and PeakForce control mechanism were used and the force-curves measured for means of set-point control in the PeakForce system and analysed in real-time to provide quantitative nanomechanical mapping (QNM) of the samples. QNM analysis was conducted with both Nusense SCOUT cantilevers [NuNano, Bristol, UK], (nominal tip radius 5 nm, spring constants 21 – 42 N m-1) and RTESPA-300 cantilevers [Bruker, CA, USA], (nominal tip radius 8 nm and spring constants 30 – 60 N m-1), whilst high resolution imaging of topographic features was conducted using SCANASYST-AIR-HR cantilevers [Bruker, CA, USA] (nominal tip radius of 2 nm). The system was calibrated for measurement of Young’s modulus (YM) fitting with DMT models, using the relative method and samples of known YM (highly oriented pyrolytic graphite (HOPG) (18 GPa) and PDMS-SOFT-1-12M (2.5 MPa) [Bruker, CA, USA]), for bone and cartilage measurements respectively. Bone was investigated in ambient environment whilst cartilage was maintained in a hydrated state post-dissection to minimise structural changes from drying. A root-mean-square (RMS) mean was calculated for 65536 measurements taken over a 500 nm x 500 nm region, 3 repeats were performed per sample; repeated on 3 fish per genotype.

### Finite Element (FE) models

Single specimens that were representative of the confocal z-stacks of 7dpf wt or mutant larvae dataset and their relative morphology were selected for the meshes. Cartilage elements were segmented in Scan IP using Otsu segmentation (Supplemental Figure 2A), then a solid geometry created using the interpolation and 3D wrap tool. Smoothing filters (recursive Gaussian at 1px3) were used on the meshes to blend any rough small element clusters.

**Figure 2:**
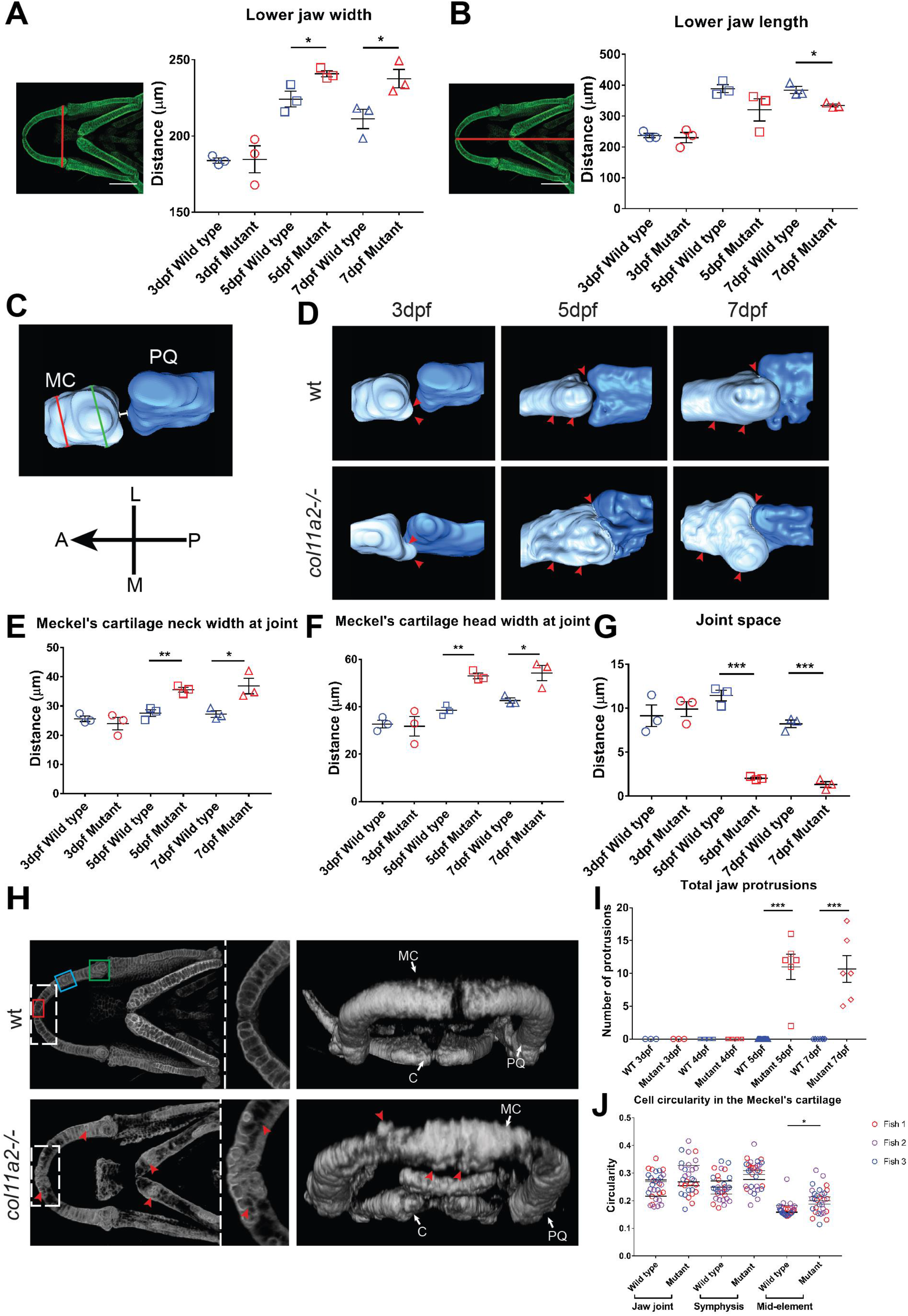
*col11a2* mutant zebrafish develop altered morphology and joint spacing in the lower jaw. A, B) Lower jaw shape quantification (n = 3 for all), location of measurements shown to the left of graphs. C) Representation of measurements taken of joint neck (red line), joint head (green line) and joint space (white line) (Meckel’s cartilage = light blue, palatoquadrate = dark blue). Orientation compass: A = anterior, L = lateral, M = medial, P = posterior. D) 3D surface renders of jaw joint from confocal images of wt and *col11a2 -/-* at 3, 5 and 7dpf. Red arrowheads = areas of change. E - G) Quantification of joint morphology at the Meckel’s cartilage neck at joint (E), Meckel’s cartilage head at joint (F) and joint space (G) (n = 3 for all). H) 3D volume renders of wt and *col11a2-/-* zebrafish at 7dpf. Dashed insets show Meckel’s symphysis at higher magnification (red arrowheads = protruding cells). I) Quantification of protruding cells in wt and *col11a2-/-* zebrafish at 3 - 7dpf (n = 3, 3, 4, 4, 13, 6, 8, 6). J) Quantification of cell circularity in the Meckel’s cartilage in 5dpf wt and *col11a2-/-* fish (n=3 for all). Location of measurements shown in H (red box = Meckel’s symphysis, blue box = mid element, green box = jaw joint). Student’s unpaired t-tests performed in A, B, E - I and J: data is mean with SEM (J shows mean with no SEM, t-tests performed between mean values). *P<0.05 P**<0.01 P***<0.001

Cartilage geometry close to the joint was separated from the main cartilage in a duplicate mesh using the 3D editing tool allowing us to assign different material properties to hypertrophic chondrocytes and immature chondrocytes (Supplemental Figure 2B). The mesh of the cartilage near the joints was subtracted from the original cartilage mesh using a Boolean operation. The meshes were added to a model and each part assigned their respective elastic isotropic material properties based on AFM measurements; values in Supplemental Figure 2, Table 1.

The models were imported into Abaqus and 2 steps created: 1 to simulate jaw closure and 2 for jaw opening. Boundary conditions were applied to these steps, with the jaw constrained in all axes of motion at the ceratohyal to anchor it in space, and in y and z at the base of the palatoquadrate. Muscle forces, direction of opening/closure and muscle attachment points were as previously described (43). The datum tool in Abaqus was used to create a custom rectangular datum coordinate system for each muscle; then used as the coordinate system for force direction between each muscle’s insertion and origin to ensure force travelled along the same vector from one end to the other. A job was created and executed for the model, and the output analysed for stress, strain and displacement.

### Measurement of jaw displacement and movement frequency

High-speed movies were made of jaw movements in wt and *col11a2* mutants; frames corresponding to maximum jaw displacements were selected, imported into ImageJ and the difference, in μm, between resting and open states at points shown in Figure 5A recorded. The number of mouth movements in 1000 frames was recorded from 7 wild type (wt) and 7 *col11a2* mutant fish as previously described in (30).

### Micro-Computed Tomography (hCT)

Three *col11a2 +/-* and three wt adult fish of the same age (1 year old) were fixed in 4% PFA for 1 week followed by sequential dehydration to 70% ethanol. Heads were scanned using an XT H 225ST micro-CT scanner (Nikon) with a voxel size of 5 μm, x-ray source of 130 kV, 53 μA and without additional filters. Images were reconstructed using CT Pro 3D software (Nikon).

### Histology

Three 1 year old *col11a2 +/-* and three wild types, were decalcified in 1M EDTA solution for 20 days. Samples were dehydrated in ethanol, embedded in paraffin and sagittally sectioned at 8 μm, relevant joint sections were de-waxed and stained with 1% Alcian blue 8GX (pH 2.5, then counterstained with Haematoxylin and Eosin. We adapted the OARSI cartilage OA histopathology grading system (44) to grade severity of OA. Five sections per jaw joint (per fish n=3 fish) were scored. PicroSirius red staining was performed using 0.1% Sirius red F3B in saturated aqueous Picric acid, washed in acidified water, dehydrated and mounted under coverslips with DPX, then imaged using polarising filters.

### Second Harmonic Generation (SHG)

SHG images were acquired from histological sections of wt and *col11a2 +/-* (n=3 fish for each genotype) using 25x 0.3 NA water dipping lens, 880nm laser excitation and simultaneous forward and backward detection (440/20) in Leica SP8 AOBS confocal laser scanning microscope attached to a Leica DM6000 upright epifluorescence microscope with multiphoton lasers and confocal lasers allowing fluorescent and SHG acquisition of the same sample and z-stack. Microscope parameters for SHG acquisition were set as described previously (45). Maximum projection pictures were assembled using LAS AF Lite software (Leica).

## Results

### *col11a2* and *col2a1* are co-expressed in the zebrafish lower jaw

To establish the extent of *col11a2* expression in cartilage we performed *in situ* hybridisation in larval zebrafish. Strong *col11a2* expression could be seen throughout the craniofacial cartilages including the Meckel’s cartilage, palatoquadrate, ceratohyal and ethmoid plate (Supplementary Figure 1A). At 3dpf, the expression pattern of *col11a2* largely overlapped the expression of the Type II collagen gene *col2a1a* visualised with the *Tg(col2a1aBAC:mCherry)* reporter zebrafish (Supplementary Figure 1B). The domain of *col11a2* expression labelled more of the joint than was labelled by the *col2a1a* transgene, and expression of both *col11a2* and *col2a1a* preceded that of the mature Type II protein, visualised by immunostaining, such that immature cells at the jaw joint and Meckel’s symphysis express *col11a2* and *col2a1a* RNA at 3dpf but not the mature Type II protein (Supplementary Figure 1C).

### *col11a2* mutants show atypical Type II collagen localisation as they develop

As Type XI collagen has previously been reported in the core of Type II collagen fibrils (3) and is thought to have a role in the stability of Type II collagen (46), we wanted to test whether loss of *col11a2* in zebrafish would impact Type II collagen stability. For this we studied the *col11a2 ^sa18324^* mutant which carries a nonsense mutation that introduces a premature stop codon at amino acid 228 (of 1877). We observed non-sense mediated decay in mutants *in situ* hybridised for the *col11a2* probe (data not shown), therefore it represents a null mutant. This mutant was crossed with the *Tg(col2a1a:mCherry)* to visualise expression of *col2a1a* and we studied its expression in craniofacial cartilages from 3-7dpf. We saw no differences in the position, timing or extent of *col2a1* expression between mutants and their siblings at 3dpf suggesting that loss of *col11a2* has no impact on the expression of *col2a1a,* although alterations to craniofacial skeletal shape in the mutant were detectable from 5dpf (Supplementary Figure 2). We next used immunostaining to detect Type II collagen protein in mutant and wild type (wt) larvae. At 3dpf we could not detect any differences between wt and mutant larvae (Figure 1A, B). However, by 5dpf clear differences in the distribution of Type II collagen were seen in the lower jaw (denoted by asterisks in Figure 1A, B). In wt fish, Type II collagen can be seen in the extracellular matrix surrounding each chondrocyte in the lower jaw cartilages, whereas in mutants, protein expression is concentrated in the perichondrium and reduced between chondrocytes (dashed insets in Figure 1B). Alongside this reduction of Type II collagen in the more mature matrix towards the middle of the cartilage elements, small pieces of immunostained material were seen separate from the main elements (red asterisks in Figure 1B) and ectopic expression of Type II collagen in the ligament was observed (red arrow in Figure 1B). Changes to the shape and size of the cartilage elements also became apparent at 5dpf, with mutant larvae displaying thicker, shorter cartilage structures, with less definition. At 7dpf these shape discrepancies were maintained. A pronounced reduction and disorganisation of Type II collagen became clear by 7dpf (Figure 1B) and was more obvious in the cartilages that make up the lower jaw, with the lateral cartilage of the ear more preserved (Figure 1A, B). Taken together these data suggest that expression and synthesis of *col2a1a* are unaffected by loss of *col11a2,* but that maintenance of Type II collagen protein is impaired in the mutants.

### Zebrafish with a *col11a2* mutation have altered jaw and joint morphology during development

Humans carrying mutations in Type XI collagen show alterations to craniofacial shape, including midface hypoplasia and micrognathia (47,48). From Type II collagen immunostaining and alizarin red/alcian blue staining, we observed that mutant zebrafish also show altered craniofacial morphology (Figures 1B and 3D). At 3dpf there was no significant difference in jaw morphology, but at 5 and 7dpf mutants had significantly shortened, wider jaws (Figure 2A, B), in line with the broader, flatter face shape observed in patients with Stickler syndrome.

As people with mutations in Type XI collagens also display abnormal joint shape (49) and increased susceptibility to osteoarthritis (8), we generated 3D surface models of the joint between the Meckel’s cartilage and the palatoquadrate (jaw joint) at 3, 5 and 7dpf in wt and mutants (Figure 2C, D). At 3dpf, there was no significant change to joint morphology, however at 5 and 7dpf, mutants show enlarged terminal regions of the skeletal rudiments. Specifically, these changes were seen at the Meckel’s cartilage neck and head adjacent to the joint (Figure 2E, F, position of measurements shown by red and green lines in Figure 2C). A reduction in the joint space (Figure 2G, position of measurement shown by white line in Figure 2C), such that the interzone is no longer clearly defined in the renders was also observed, likely due to the increased local deposition of Type II collagen (Figure 1B and 2D). These results show that the abnormal pattern of Type II collagen deposition seen in *col11a2* zebrafish mutants at 5 and 7dpf leads to altered joint shape.

### Mutation of *col11a2* leads to altered chondrocyte cell behaviour

Using confocal imaging of a transgenic reporter for *col2a1a* we were also able to observe alterations to chondrocyte behaviour in mutant fish at 7dpf. In wt larvae all cells expressing *col2a1a* are located within the cartilage element, however, in mutants we frequently observed chondrocytes located outside the main body of the cartilage element (Figure 2H). We quantified the number of these cells in the lower jaw cartilages of wt and mutant larvae during early development. Prior to 5dpf these spurs, which resemble hereditary multiple exostoses (50), are not observed (Figure 2I). However, at 5dpf and 7dpf they are present in the Meckel’s cartilage and ceratohyal of *col11a2* mutants, suggesting a failure of chondrocyte progenitor cells to fully intercalate into the cartilage element prior to expression of *col2a1a.*

In addition to these protrusions, we observed differences to chondrocyte morphology within the jaw elements of mutants at 5dpf. Chondrocyte circularity is a measure of maturation, as in zebrafish chondrocytes become less rounded over time and form tightly packed stacks as they mature towards hypertrophy and show organisation reminiscent of the mammalian cartilage growth plate. Chondrocytes in the centre of cartilage elements (representing the most mature chondrocytes) of *col11a2* mutants show increased circularity over those in wt cartilage (Figure 2J). These results suggest that the maturation of chondrocytes in *col11a2* mutants is disrupted, potentially due to the loss of Type II collagen from the ECM, which could provide an explanation for the thicker appearance of the cartilage elements in mutants.

### Larval and adult *col11a2* mutant zebrafish have altered material properties in the craniofacial skeleton

To test whether loss of Type II collagen and alterations to chondrocyte behaviour led to changes to the material properties of the cartilage, we performed atomic force microscopy (AFM) on dissected lower jaw cartilages from 7dpf wt and mutants. We tested the properties in regions containing immature cells close to the jaw joint and Meckel’s symphysis (Figure 3A) and observed a significant increase in Young’s modulus (YM) from an average of 4.15 to 7.4 MPa (Figure 3B). In more mature, intercalated cells towards the centre of the Meckel’s cartilage (in which we saw loss of Type II collagen) the difference in YM was around 4 times greater than that of comparable regions in wt (Figure 3B). This suggests that loss of Type II collagen as a result of the *col11a2* mutation leads to stiffening of the cartilage ECM. As we observed altered material properties in larvae at pre-skeletonised stages we wanted to test whether this would persist to adulthood and impact bone properties. We dissected jaw bones and operculae from wt and mutant fish and used AFM to establish YM. As in the larval cartilage, we observed that the bone from *col11a2* mutant fish had a significantly higher YM than siblings (Figure 3C).

**Figure 3:**
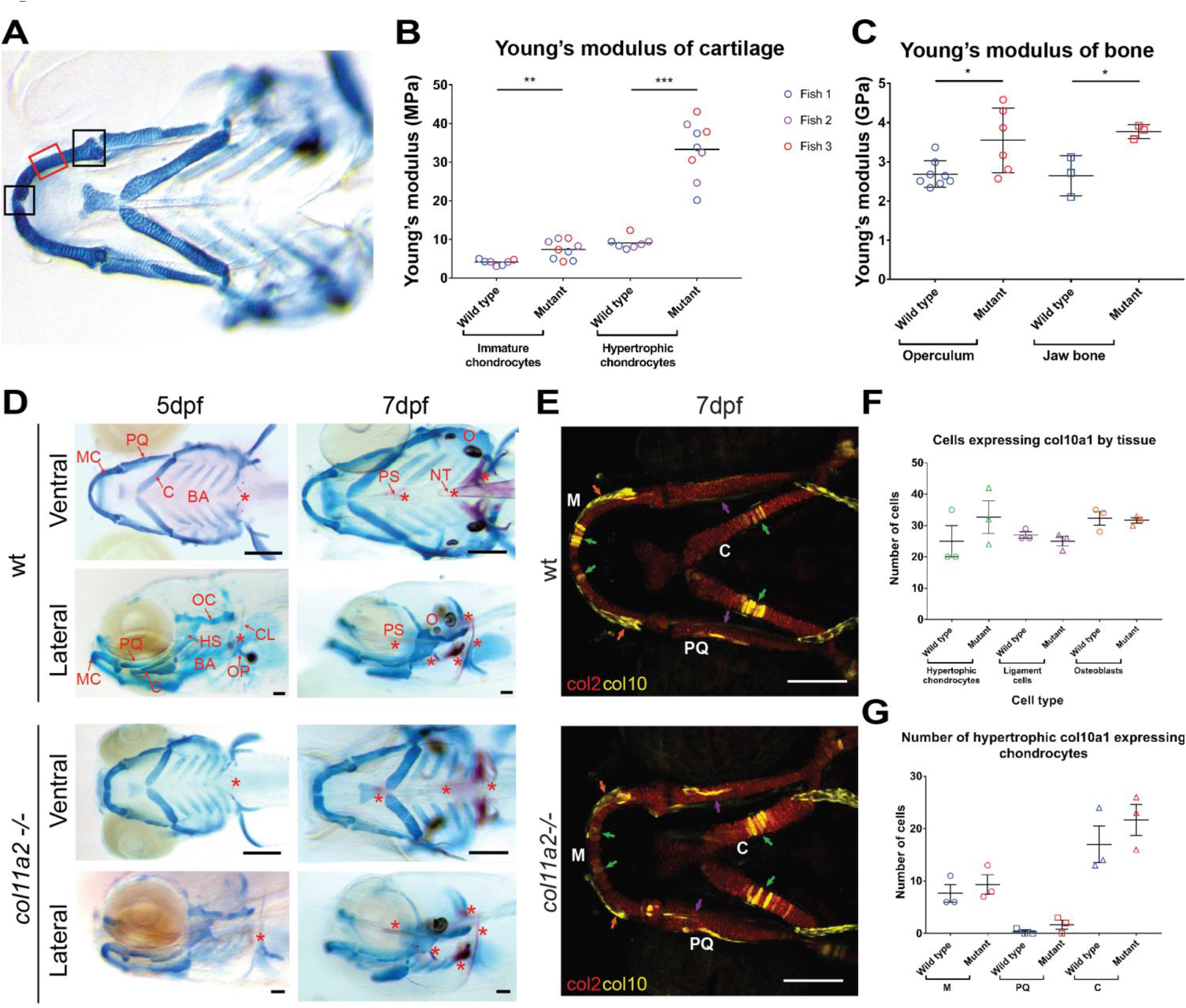
*col11a2* mutants have altered material properties in more mature cartilage which is not explained by increased mineralisation or hypertrophy. Location of AFM measurements taken from larvae shown in (A). Measurements for immature chondrocytes taken from either of the two areas marked by black boxes, measurements for hypertrophic chondrocytes taken from area marked by red box. B, C) Young’s modulus values for (B) immature and hypertrophic chondrocytes in wt and *col11a2-/-* (n=3 for both) at 7dpf and (C) adult bone from the operculum and jaw in wt (n=8 and 3 respectively) and *col11a2-/-* (n=6 and 3 respectively). D) Ventral and lateral views of alizarin red alcian blue staining show glycosaminoglycans in cartilage (stained in blue) and mineralisation (stained in red) in wt and *col11a2-/-* fish at 5 and 7dpf. Red asterisks indicate areas of bone formation. MC = Meckel’s cartilage, PQ = palatoquadrate, C = ceratohyal, BA = branchial arches, HS = hyosymplectic, OC = otic capsule, OP = operculum, CL = cleithrum, PS = parasphenoid, NT = notochord tip, O = otoliths. Scale bar = 200μm. E) *col10a1aBAC:citrine;col2:mCherry* transgenic line shows Type X (yellow) and Type II (red) collagen in wt and *col11a2-/-* zebrafish at 7dpf. Scale bar = 100μm. F) Quantification of col10a1 expressing cells in hypertrophic chondrocytes, IOM ligament cells and osteoblasts in the lower jaw at 7dpf (position of each cell type shown by green, purple and orange arrows in (E) respectively) (n=3 for all). G) Quantification of *col10a1* expressing hypertrophic chondrocytes in 7dpf wt and *col11a2-/-* fish (n = 3 for all) (M = Meckel’s cartilage, PQ - palatoquadrate, C = ceratohyal). Student’s unpaired t-tests were performed in B, C, F and G, data is mean with SEM (B shows mean with no SEM). *P<0.05 P**<0.01 P***<0.001

### Type II collagen loss is not accompanied by changes to GAGs, Type X or Type I collagen in *col11a2* mutant zebrafish

As Type II Collagen was prematurely lost from maturing chondrocytes, we sought to examine whether glycosaminoglycans (GAG)s, were similarly reduced. We stained wt and mutant larvae with Alcian blue (to mark cartilage GAGs) and Alizarin red (to mark bone) at 5 and 7dpf. We saw no reduction of GAG reactivity in the mutants at 5 or 7dpf (Figure 3D). Additionally, we saw no dramatic changes to Alizarin red, with dermal and chondral bones in mutants mineralising at a similar rate to wt fish.

During cartilage maturation, a change in collagens from Type II to Type X collagen is associated with chondrocyte hypertrophy (51). In teleosts such as zebrafish *col10a1a* marks hypertrophic chondrocytes, but also osteoblasts and ligament cells (52,53). To test whether there was any change to the extent of chondrocyte hypertrophy or to the number of osteoblasts or skeletal connective tissue cells we crossed the *col11a2* mutant into the *col10a1a* transgenic reporter (Figure 3E). Quantification of *col10a1a* positive cells in wild type and mutant fish revealed no differences to the number of hypertrophic chondrocytes at 7dpf (Fig 3F green arrows and graphs Fig 3.G,H), we also saw no difference in the number of osteoblasts in the dentary which is located directly adjacent to the MC (red arrows Fig 3F, quantification in 3G) or to the number of cells in the IOM ligament (purple arrows in 3F, quantification in 3G). This suggests that the onset of hypertrophy is not disrupted in *col11a2* fish, despite the ‘less mature’ appearance of their chondrocytes.

During cartilage degeneration, such as in osteoarthritis, a switch of collagens is commonly reported with a reduction of Type II and increase in Type I collagen, associated with stiffer matrix (54). To test whether loss of Type II collagen led to replacement with Type I, we performed immunostaining in wt and mutant larvae at 5 and 7dpf. In wt fish Type I collagen was present in the jaw joint space, Meckel’s symphysis and at a low level in the cartilage ECM, this pattern was unchanged in mutants at 5dpf, however by 7 dpf there was a reduction of Col I in the joint interzone of mutants but no change within the cartilage elements themselves (Supplementary Figure 4).

**Figure 4:**
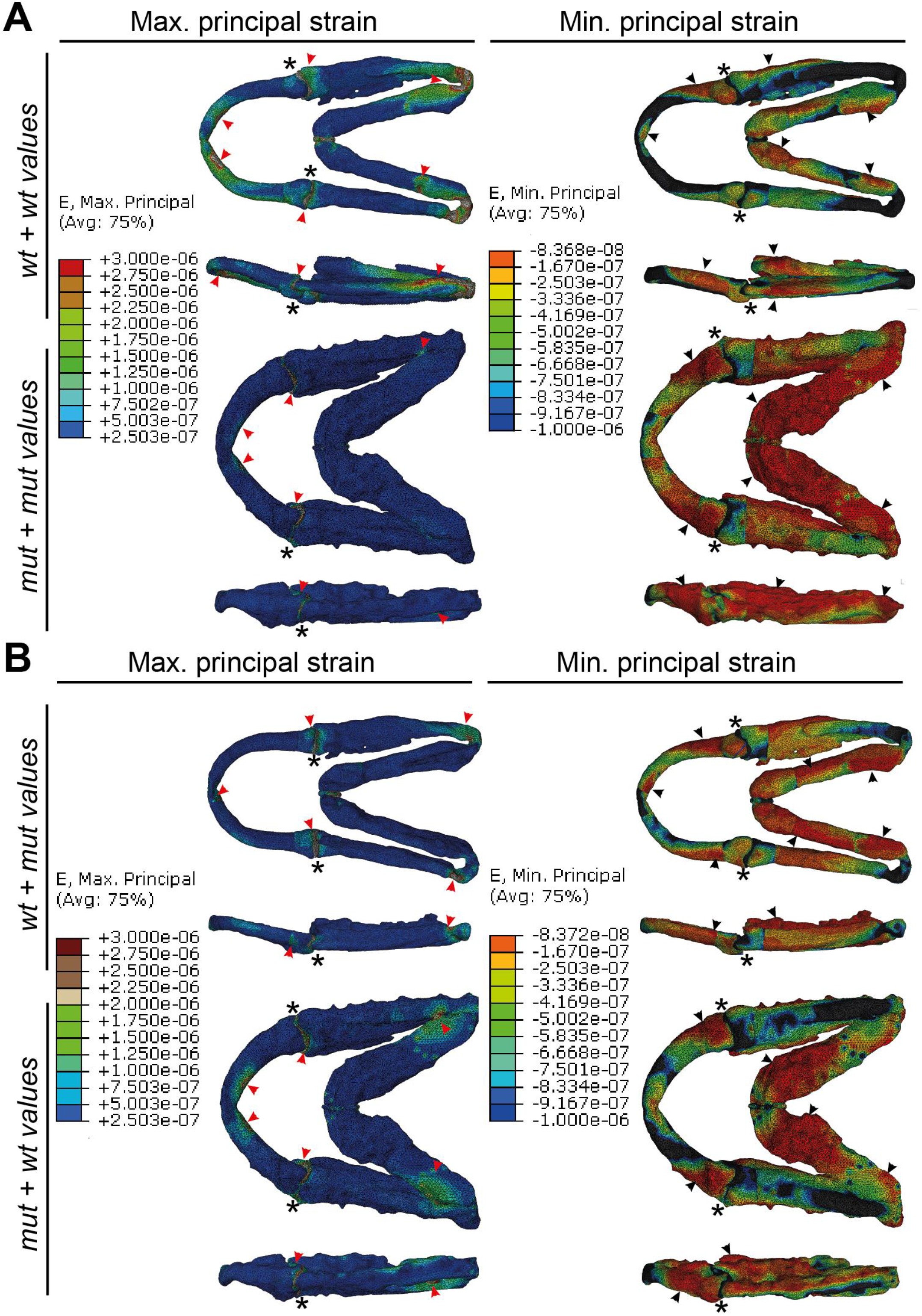
Shape changes in *col11a2* zebrafish mutants have a greater effect on jaw biomechanics than material property changes. A, B) Finite Element (FE) models of maximum (E. Max) and minimum (E. Min) principal strain during mouth opening in 7dpf wt and *col11a2-/-* zebrafish. Red arrowheads = areas of high strain, black arrowheads = areas of low strain, black asterisks = jaw joint. A) wt jaw shape with wt material properties and *col11a2-/-* shape with *col11a2-/-* material properties. B) wt shape with *col11a2-/-* material properties and *col11a2-/-* shape with wt material properties. Ventral and lateral views shown for each condition.

Taken together, these results suggest that loss of Type II collagen in *col11a2* mutants does not lead to loss of GAG, nor to compensatory increases of Col I or *col10a1,* and that hypertrophy is unaffected by the *col11a2* mutation. As a result, the increase in cartilage stiffness observed from AFM cannot be attributed to alterations in these components.

### Both shape and material properties impact the biomechanical performance of the zebrafish lower jaw

We have previously modelled the biomechanics of zebrafish jaw opening and closure during early ontogeny using Finite Element Analysis (FEA) (43) and shown that paralysis and the accompanying changes to joint shape impact the strain pattern in the developing joint (33). Therefore, we used FEA to model how the changes to shape and material properties observed in *col11a2* mutants would affect the biomechanical performance of the lower jaw. Meshes were generated of wt and mutant larvae at 7dpf (meshes shown in Supplementary Figure 5B). We applied muscle forces as per Roddy et al (55) and used the material properties established from AFM (Figure 3B).

**Figure 5:**
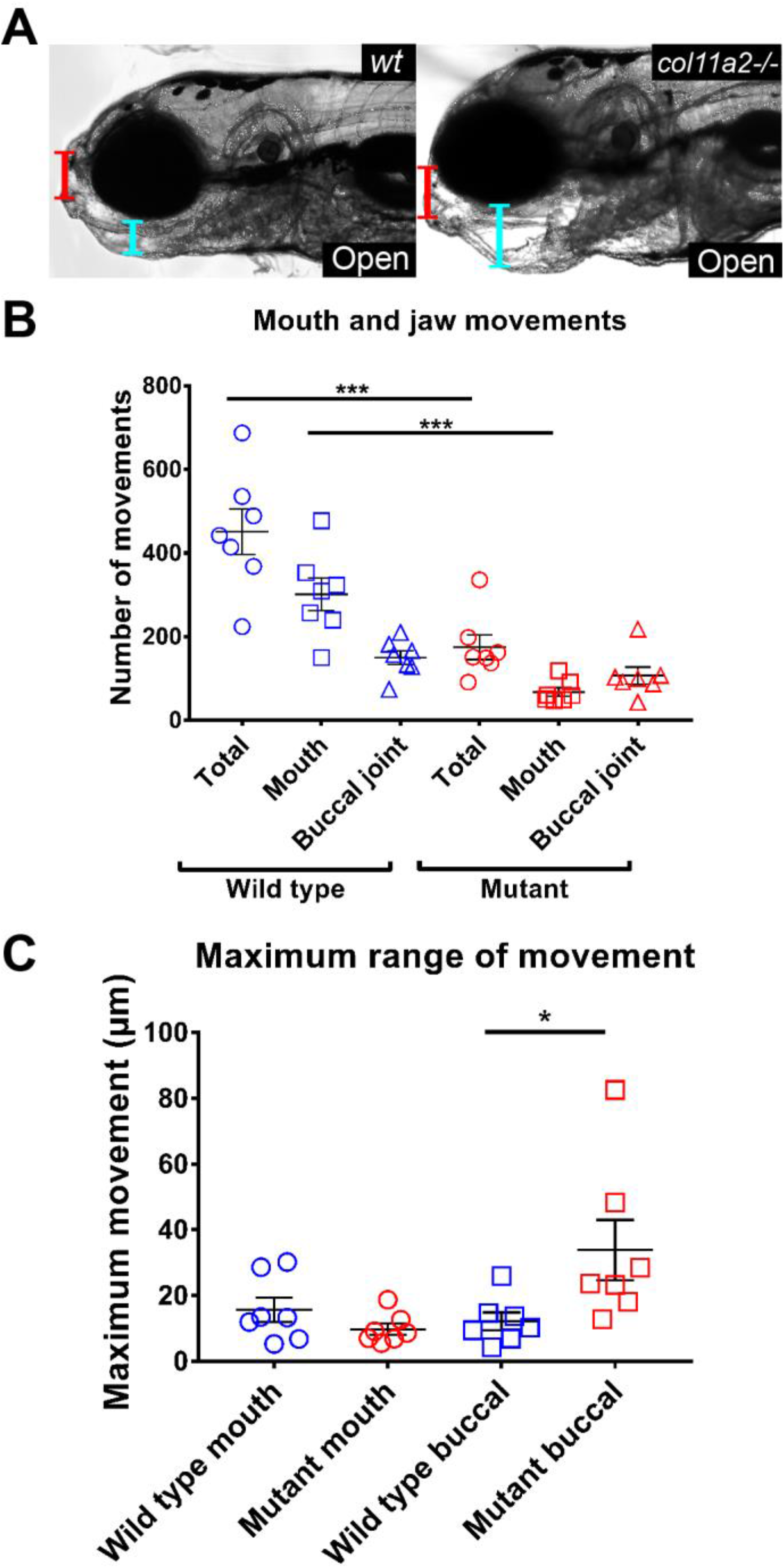
*col11a2* mutant zebrafish have abnormal jaw movement at 5dpf. A) Stills from high speed movies show range of jaw movement in wt and *col11a2-/-* B, C) Analysis of (B) total jaw movements and (C) range of movement at two locations shown in (A): red line = mouth, blue line= buccal joint (n= 7 for all). Student’s unpaired t-tests performed for B and C, data is mean with SEM. *P<0.05 ***P<0.001.

We first modelled the wt and the mutant jaw shapes using the material properties established from each genotype and modelled a 2 step process for jaw movement with Step 1 denoting jaw closure and Step 2 jaw opening (as per (43,55)) and visualised the maximum principal (EMax) and minimum principal (EMin) strains for jaw opening (Figure 4) and closure (Supplementary figure 6). In the wt model, tensional strains (EMax) are located laterally around the joint and either side of the Meckel’s symphysis, focused on the muscle insertion points with the strain spreading widely through the element. By contrast, in mutants, maximum principal strain is concentrated on the joint interzone, with little spread through the cartilaginous element (note blue colour throughout the cartilage of mutants, c.f. greens and yellows in wt) (Figure 4A, Supplementary videos 1,2). Comparison of the Meckel’s symphysis joint interzones for strains (EMax) showed that the difference between the types of models (wt models and the mutant models) was far larger than the difference between each type of model with different material properties (Supplementary Figure 5C, D). In the mutant model with mutant values the joints averaged 1.63E-02 maximum strains (EMax), and in the mutant model with wt values the joint strain averaged 1.51 E-02, whereas in the wt models, the joint strains averaged 4.41E-02 (EMax) in the wt with wt values model, and 5.14E-06 in the wt with mutant values model. Between each type of model (wt against mutant), the average maximum principal strains have not changed substantially despite change in material properties. In wt, compressional strains (EMin) are at the Meckel’s symphysis, the medial surface of the anterior MC and on the dorsolateral side of the jaw joint (Figure 4A, Supplementary Figure 6A). In mutants, again, the minimum principal strains are more focal than in wt larvae (Figure 4A, Supplementary Figure 6A).

Changing the material properties of the models affects patterns of strain and displacement. This influences the displacement on the overall jaw morphology as seen in the jaw opening stage when both models are given the stiffer mutant material properties. In these models, the jaw shows less displacement in the opening movement compared to the situation in which models are given the less stiff wildtype material property values.

To test whether the change to the strain pattern was predominantly caused by the alteration to the shape of the jaw elements or the changes to YM, we next modelled the effect of mutant properties in the wt shape, and wt properties in the mutant shape (Figure 4B, Supplementary Figure 6B). We observed that changing YM in the wt shape to the mutant values decreased the spread of max and min principal strains, such that the pattern was intermediate between the mutant and wt. Likewise, inserting the wt values for cartilage into the mutant shape lead to an increase in the extent of both tension (EMax) and compression (EMin). However, it did not fully ‘rescue’ the pattern, leading us to conclude that while both shape and material properties play a role in the mechanical performance of the tissue, the effect of shape is greater than that of material properties.

### *col11a2* zebrafish mutants show impaired jaw function

As patients with Stickler syndrome suffer from joint hypermobility (56), and as *col11a2* zebrafish mutants show aberrant joint morphology, we looked at jaw function at 5dpf. Zebrafish have two joints within the lower jaw and make distinct movements for feeding and breathing (57). By taking movies and quantifying jaw movement, we observed that mutants make significantly fewer total movements than wt (Figure 5B, Supplementary videos 3,4). This was due to a reduction in the number of movements involving the jaw joint, as we observed no significant difference in the frequency of movements involving the buccal joint (Figure 5B). However, mutant zebrafish show an increased range of motion at the buccal joint, which appears to dislocate (Figure 5A, C). To rule out the possibility that this change to movement was caused by altered muscle patterning, we quantified the number of slow twitch fibres in the jaw at 5dpf and saw no difference in fibre number between wt and mutants (Supplementary Figure 7A,B). We also measured the diameter of the intermandibularis posterior and interhyoideus muscles in the lower jaw from birefringence and found no change in diameter between wt and mutants (Supplementary Figure 7C). Taken together this suggests that the alterations to joint shape observed in *col11a2* mutants are the cause of abnormal joint function.

### Premature OA is observed in adult *col11a2* heterozygous fish

Due to the abnormalities in joint shape, mechanical performance and function in mutants, and since aberrant joint loading is highly associated with OA risk (58), we wanted to test whether adult mutants would develop premature osteoarthritis. To address this question, we analysed 1-year old heterozygous fish *(col11a2+/-)* and wt siblings using micro-computed tomography (μBC;CT). Craniofacial abnormalities were observed in *col11a2+/-,* including jaw protrusion and hypoplasia of the fronto-nasal bone (Figure 6A). Changes in joint shape were observed in *col11a2+/-* accompanied by narrowing of the inter-joint space (Figure 6B). To identify the histopathological changes related to OA, we stained wt and *col11a2+/-* joint sections for Alcian Blue and H&E (Figure 6C). While in wt sections a defined cartilaginous layer lines the joint, in *col11a2+/-* the cartilage shows signs of degradation. Grading of 5 sections per joint per fish (n=3 fish) using the criteria in the OARSI scoring system (44) showed an average score of 6 in the *col11a2+/-* sections which is characterised by deformation and change in the contour of the articular surface, compared to an intact surface and normal cartilage with average score of 0 in the siblings (Figure 6C). Osteophytes were not observed. We analysed collagen organisation using PicroSirius red staining and Second Harmonic Generation (SHG) (Figure 6D and E). In wt jaws, the cartilaginous layer at the joint shows organised collagen fibres with a distinct orientation from those of the underlying bone (Figure 6D, note change in colour from red to green in PicroSirius red staining). However, in the *col11a2+/-* the transition from cartilage to bone is lost and the overall organization is perturbed (Figure 6D). Thicker collagen bundles and fibres displaying abnormal orientations were seen through SHG in *col11a2+/-* samples (Figure 6E). Taken together these data demonstrate that loss of *col11a2* leads to early onset of osteoarthritis-like changes in adults.

**Figure 6:**
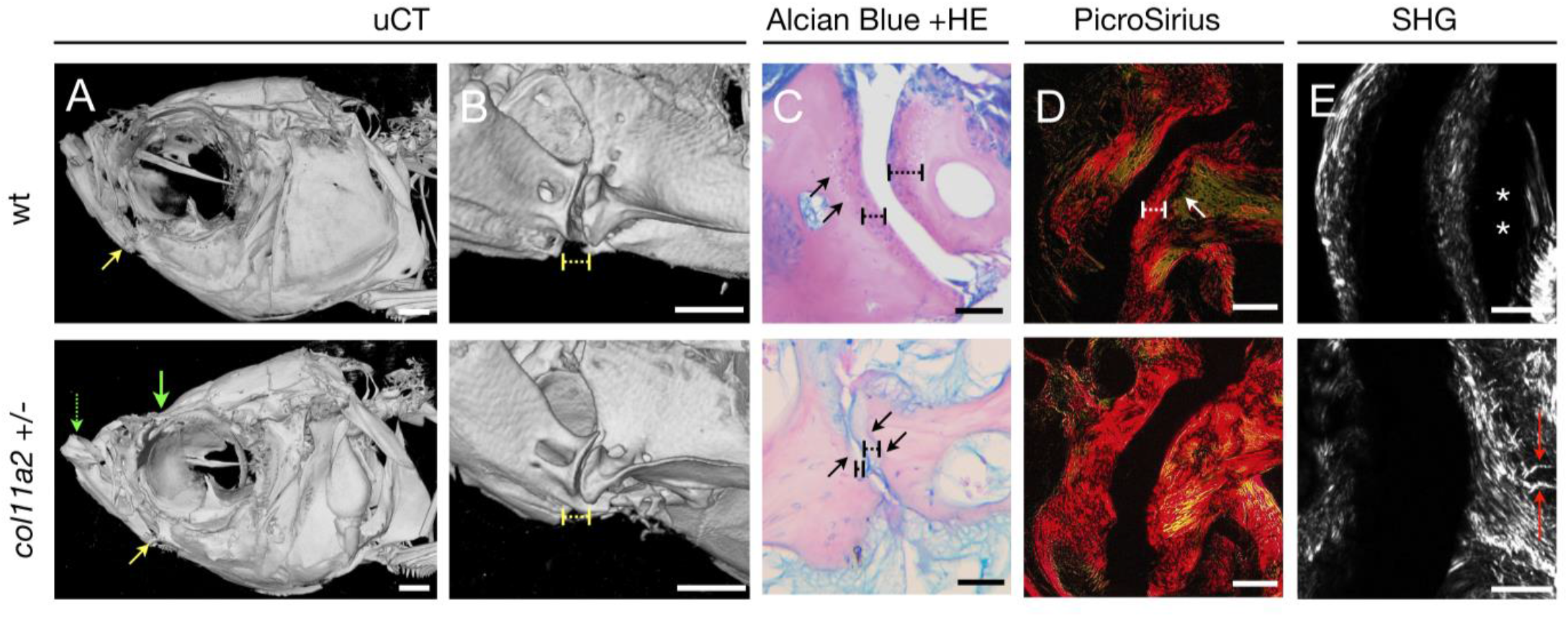
Mutations in zebrafish *col11a2* result in changes that trigger premature OA. A, B) 3D renders from μCTs of 1-year old wt and *col11a2* heterozygous mutant *(col11a2+/-).* A) Yellow arrow = jaw joint, dashed green arrow = region of jaw protrusion in *col11a2+/-,* green arrow = region of hypoplasia in fronto-nasal skeleton. B) Higher magnification image of joint region where dashed yellow line = inter-joint space. C, D) Paraffin sections of the jaw joint stained with (C) Alcian blue and Haematoxylin/Eosin and (D) PicroSirius red. Dashed black line = cartilage layer, black arrows = underlying bone, dashed white line = cartilage, white arrow = bone (green). E) Second Harmonic Generation (SHG), asterisks pointing to areas of thinner fibres not detected by SHG, Red arrows = thicker collagen bundles on abnormal orientation. Scale bars = 50μm.

## Discussion

Mutations in the Type XI collagen genes *col11a1* and *col11a2* have previously been linked to numerous skeletal dysplasias, such as Stickler syndrome and Fibrochondrogenesis, which are associated with cartilage destabilisation, and abnormal skeletal shape and properties. Here, we describe the impact of loss of *col11a2* in zebrafish and show changes to ECM composition, material properties, craniofacial shape, mechanical performance, chondrocyte behaviour, and joint function in larval and adult fish.

Type XI collagen is important for the protection of Type II collagen from degradation (59); our data suggest that, while transcription and secretion of Type II collagen is unaffected at early stages of larval development, the assembly of Type II collagen fibrils may be altered in mutants, making them more susceptible to degradation. This idea is given weight by the identification of fragments of Type II positive material seen surrounding the cartilage elements. What happens to those degraded collagen fragments is still unclear. Potentially, they may be cleared by the phagocytic cells of the innate immune system either with a rapid resolution, or alternatively, continued accumulation of these fragments could lead to the low-level inflammation associated with osteoarthritis (60,61). Loss or breakdown of Type II collagen also occurs as the chondrocytes mature, such that the matrix between the chondrocytes almost completely lacks Type II collagen, while the matrix of the perichondrium is relatively preserved. We have tested effects of *col11a2* loss on the material properties of cartilage and bone, and our data shows an increase in YM in both tissues, with the greatest difference seen in mature chondrocytes. It may be noted that Young’s modulus for zebrafish cartilage is higher than that from other species (4.15MPa in fish vs 0.45Mpa in human articular cartilage (62)). One likely explanation is the variation in relative ratio of cells to matrix during development and across species in evolution. In mature human articular cartilage the ratio is approximately 10:90 cells to matrix compared to 80:20 in zebrafish (63). The higher YM in mutants was not explained by any obvious increased calcification, accumulation of Type X or Type I collagen or loss of GAG. In rodent models increased matrix stiffness has been described as chondrocytes mature in the growth plate (64,65). The stiffness of collagen matrix is controlled by several factors including fibre diameter and the density of intra-fibrillar cross-links, and abnormalities in collagen fibrillar assembly have been related to changes to mechanical properties of the cartilage during progression of OA (66) and ageing (67).

It has previously been reported that patients with Stickler syndrome develop premature OA (1) but the mechanism by which this occurs is unclear. We and others have previously shown that, despite living in an aquatic environment, zebrafish can also develop alterations to the joint that strongly resemble OA (68,69). Interestingly, we see premature development of osteoarthritis in *col11a2* heterozygous adult zebrafish. This is manifested by abnormal collagen organisation, degeneration of joint cartilage and loss of joint space. During OA, proteoglycans are lost from the cartilage prior to the degradation of the collagen network in the extracellular matrix (2). This change to the organisation and content of collagen in the cartilage leads to changes in its material properties (3), including its stiffness and tensile strength (4). It has previously been demonstrated that in OA, cartilage stiffness is often reduced (17,70)while we saw a dramatic increase in cartilage matrix stiffness in the *col11a2* mutants these measurements were taken from larvae. We saw increased YM in adult bone, both of dermal (operculum) and chondral (jaw) bone, albeit less dramatically than in the cartilage. Potentially, stiffer bone could exacerbate OA pathogenesis; as subchondral bone thickening accelerates the degradation of articular cartilage (71). Alternatively, and perhaps more likely, changes to joint loading from the abnormal shape and function throughout life may be the driver for the development of pathogenic osteoarthritis-like changes in the joint. To test this more fully it would be desirable to follow the development of the pathology throughout the life course of the fish.

As joint mechanical performance is impacted by its shape and the material properties of the tissues, we explored the relative impact of each by testing the impact of altering material properties in the wild type and mutant shapes. From this we deduced that while both contribute to the strain pattern, the larger impact comes from joint architecture. However, questions remain to the exact sequence of events; are the increases in YM in immature chondrocytes sufficient to drive local changes to cell behaviour within the joint? If so subtle changes to joint morphology, could impact joint mechanics upon onset of function, leading to further, more significant changes to skeletal cell behaviour. Movement of joints has been shown to be required for their correct specification in the majority of joint types in all species studied (72–74,26,75). Interestingly, at the earliest stages we studied (3dpf), prior to the onset of joint movement, the mutants are barely distinguishable from wild types, despite the *col11a2* gene being expressed throughout the cartilage from 2dpf. Following the onset of movement changes between wild type and mutants become more pronounced, these include the degradation of Type II collagen from the mature matrix, and the loss of the joint space. This loss of correct joint spacing and the enlargement of the rudiments could be explained by premature differentiation of the immature cells of the interzone. A requirement for normal movement has been demonstrated in chick, mouse and fish to maintain joint space and to prevent ectopic expression of Type II collagen (33,34,76,77). Alternatively, it could represent a failure to maintain local gdf5 signalling; it has recently been shown that there is a requirement for the continued influx of Gdf5 positive cells for correct joint specification (78).

It is likely that by changing the mechanical performance of the joint, that mechanosensitive genes will be differentially activated, and these likely control the cellular changes we describe. Candidates that could be differentially activated in the mutants could include the Piezo ion channels which have been shown to play a role in OA (79). Another candidate could be the YAP pathway; YAP is implicated in negative control of chondrogenesis (80,81). Or the genes in the Wnt signalling pathway. The Wnt pathway has been implicated in developmental skeletal mechanosensation in mice, chicks (82) and zebrafish (83), and could potentially be acting in combination with BMP regulatory genes such as Smurf1 (84). We have shown in zebrafish that wnt16 controls chondrocyte proliferation and migration in the joint region. Wnt16 is also linked to hip geometry (85) altered cortical bone thickness (86,87) the response of chondrocytes to injury and to osteoarthritis (88,89).

Following the onset of movement, we also see the appearance of cells located outside the cartilage anlage, which bear some resemblance to multiple hereditary extoses (MHE). Stickler syndrome is associated with MHEs (90). It has been reported in a zebrafish model that the development of MHE is driven by changes to the matrix from loss of the Extosin genes, that, while dispensable for early chondrocyte differentiation are required for chondrocyte maturation, hypertrophy and intercalation and which encode genes lead to matrix sulfation (50). Potentially, the loss of Type II collagen in the *col11a2* mutants could perturb sulfation. Alternatively, these cells could fail to intercalate then be extruded due to altered joint function, as paralysis has been shown to control chondrocyte intercalation in zebrafish (91). The failure of these cells to fully intercalate leads to shorter, thicker elements in *col11a2* mutants.

Taken together our findings show that loss of *col11a2* in zebrafish leads to changes to matrix phenotype, and cell behaviour that impact the biomechanical and functional performance of the developing joint leading to premature OA. By making use of the detailed dynamic imaging unique to small translucent models like the zebrafish we were able to follow the alterations to the developing skeleton at cellular resolution, identifying changes to cell behaviour that go some way to explaining how loss of a relatively minor collagen subtype can have such a profound effect on the human skeleton in diseases such as Stickler syndrome and Fibrochondrogenesis.

## The authors declare no competing interests

EL, KR and EK carried out the molecular lab work, participated in data analysis, participated in the design of the study and drafted the manuscript; JA generated the FE models; RH performed AFM; CLH conceived the study, designed the study, coordinated the study and helped draft the manuscript. All authors gave final approval for publication.

## Acknowledgements

The authors would like to thank Stephen Cross and the Wolfson Biomaging facility staff for help with image acquisition and analysis.

